# First national survey of terrestrial biodiversity using airborne eDNA

**DOI:** 10.1101/2025.04.07.647580

**Authors:** Orianne Tournayre, Joanne E. Littlefair, Nina R. Garrett, James J. Allerton, Andrew S. Brown, Melania E. Cristescu, Elizabeth L. Clare

## Abstract

Near real-time data across taxa are necessary for quantifying biodiversity at regional to continental scales and evaluating conservation measures. Yet, standardized methods and globally distributed infrastructure are still lacking. In this study, we conducted the first national survey of terrestrial biodiversity using a metabarcoding approach on airborne environmental DNA collected by a national ambient air quality monitoring network. Our goal was to perform a multi-taxonomic biodiversity assessment at a national scale, compare detections with those of another large-scale monitoring approach (citizen sciences) and estimate a tentative minimum eDNA transportation distance. We identified over 1,100 taxa, including vertebrates, invertebrates, protists, fungi and plants covering a wide range of life history traits and ecological niches. Citizen science and eDNA detections were complementary, with eDNA better mapping less charismatic and difficult to spot taxa, demonstrating its potential to align with global conservation goals. Airborne eDNA signals were relatively local, likely due to the deposition of the larger particles from the air over shorter distances and limited wind transportation at near ground level. Overall, our results show that molecular protocols integrated into existing air quality monitoring networks can provide standardized, biodiversity monitoring at relatively low field cost, with potential for broad scalability.

## INTRODUCTION

Biodiversity is experiencing an accelerated decline due to habitat modification, overexploitation, invasive species, pollution, and climate change^1^. Beyond the alarming loss of species, these factors cause major shifts in biodiversity distribution and community composition. Ultimately these shifts can cascade to the disruption of ecosystem functions^2^. The severity of this trend prompted international agreements, such as the Convention on Biological Diversity, and is included in multiple UN Sustainable Development Goals (e.g. SDG 13, 14, 15). Long-term biodiversity surveys, which catalogue species presence and abundance over time at fixed locations are essential for detecting these changes^3^. The ultimate goal is to measure the dynamics of communities in real time and assess the success or failure of interventions to mitigate biodiversity loss. Achieving this requires high-resolution spatio-temporal data collected regularly across large scales and multiple taxa. While there are mechanisms to monitor some species this way, methodologies vary widely^4–6^ making traditional taxon-specific surveys too labor-intensive and impractical for broad applications.

Consequently, biodiversity data are almost always logistically and taxonomically narrow and limited in time and space^3^. The most limiting factor is the lack of infrastructure for large-scale biodiversity monitoring.

One approach to large scale monitoring is to capitalize on citizen science programs. These are a powerful tool to mitigate data limitations by leveraging the efforts of local volunteers to gather large amounts of data across vast geographic areas and extended time periods^7^.

These datasets can address many purposes, such as estimating population trends and species distributions, and have the added benefit of increasing public interest and awareness. As with any method however, these largely opportunistic datasets can be temporally (e.g. spring activity^8^), spatially (e.g. over 94% of iNaturalist observations are within 1 km of roads in British Columbia, Canada^9^), or taxonomically (e.g. easily recognizable, charismatic taxa^10^) biased^11,12^. However, they provide an unprecedented amount of data with good coverage for certain use cases, and serve as a powerful complementary monitoring tool. For instance, they can be efficiently combined with emerging technologies, such as remote sensing, artificial intelligence (AI) classification of images or acoustic data, and DNA techniques. For just a few examples, August et al.^13^ created a new biodiversity dataset from social media by using an AI image classifier on 60,000 geolocated images, and “Walrus from Space’’, which relies on users looking for wildlife on public satellite pictures, has led to the rediscovery of a beluga whale population previously thought to have been extirpated^14^.

DNA metabarcoding is a molecular approach providing the parallel identification of multiple species based on short target gene regions. It allows the massive and rapid identification of unsorted aquatic or terrestrial organisms in bulk collections^15,16^, but also more recently in environmental samples. It has been particularly well used on freshwater and marine water samples, targeting various groups of vertebrates, invertebrates, plants and phytoplankton^17^. By overcoming the need to directly capture or observe organisms to identify them, environmental DNA (eDNA) has offered new possibilities to assess the state of biodiversity on a global scale in near real time^18^. For example, the 2023-2025 Global Lake Sampling collaborative project (ETH Zürich) aims at monitoring both animal and plant aquatic diversity at a global scale by sampling water from hundreds of lakes on the same day across the planet. This represents an incredible spatial assessment but still relies on the labour of volunteers and is not temporally replicated. Vertebrate communities have most frequently been investigated using water samples (review in ^19^), while other terrestrial communities have primarily been investigated using soil (e.g. plants^20^), or fecal samples (e.g. arthropods, vertebrates, plants in generalist predator feces^21^). Airborne eDNA has recently emerged as a powerful tool for terrestrial ecology collected using either passive (e.g. dust traps^22^) or active samplers^23^. It was first used to detect bacteria, fungi and both wind-pollinated (anemophilous) and non-anemophilous plant species, but recent studies showed its applicability to other terrestrial taxonomic groups including vertebrates^24^ and arthropods^25^. As with other eDNA-based approaches, airborne eDNA detection can be affected by environmental factors, degradation, transportation, and some uncertainties in taxonomic resolution. However, it has one unique property among eDNA tools because it has the extraordinary potential for long-term observation of local biodiversity on a continental scale using the existing infrastructure of pollution monitoring networks. A proof of concept using such infrastructure on two sites over a month recovered the eDNA of 183 vertebrates, arthropods, plants and fungi as a byproduct of regular operation^26^. If expanded, such campaigns could overcome the key challenges of field labour, taxon specificity, replication and sampling scale instantly.

In this study, we demonstrate the scalability of this approach by expanding this proof of concept to conduct the first national survey of broad terrestrial diversity using airborne eDNA relying on existing air monitoring infrastructure with no modification of its operation. The UK is an ideal model as a relatively small country with extensive knowledge of biodiversity from abundant records on wildlife distribution. Our goals were to 1) assess vertebrate, arthropod, plant and fungi diversity coverage across the country using this data source (Figure 1), 2) compare these detections with one of the only monitoring approaches providing such large-scale coverage, i.e. citizen sciences records, and 3) estimate potential airborne eDNA spatial transportation distances.

**Figure 1.**
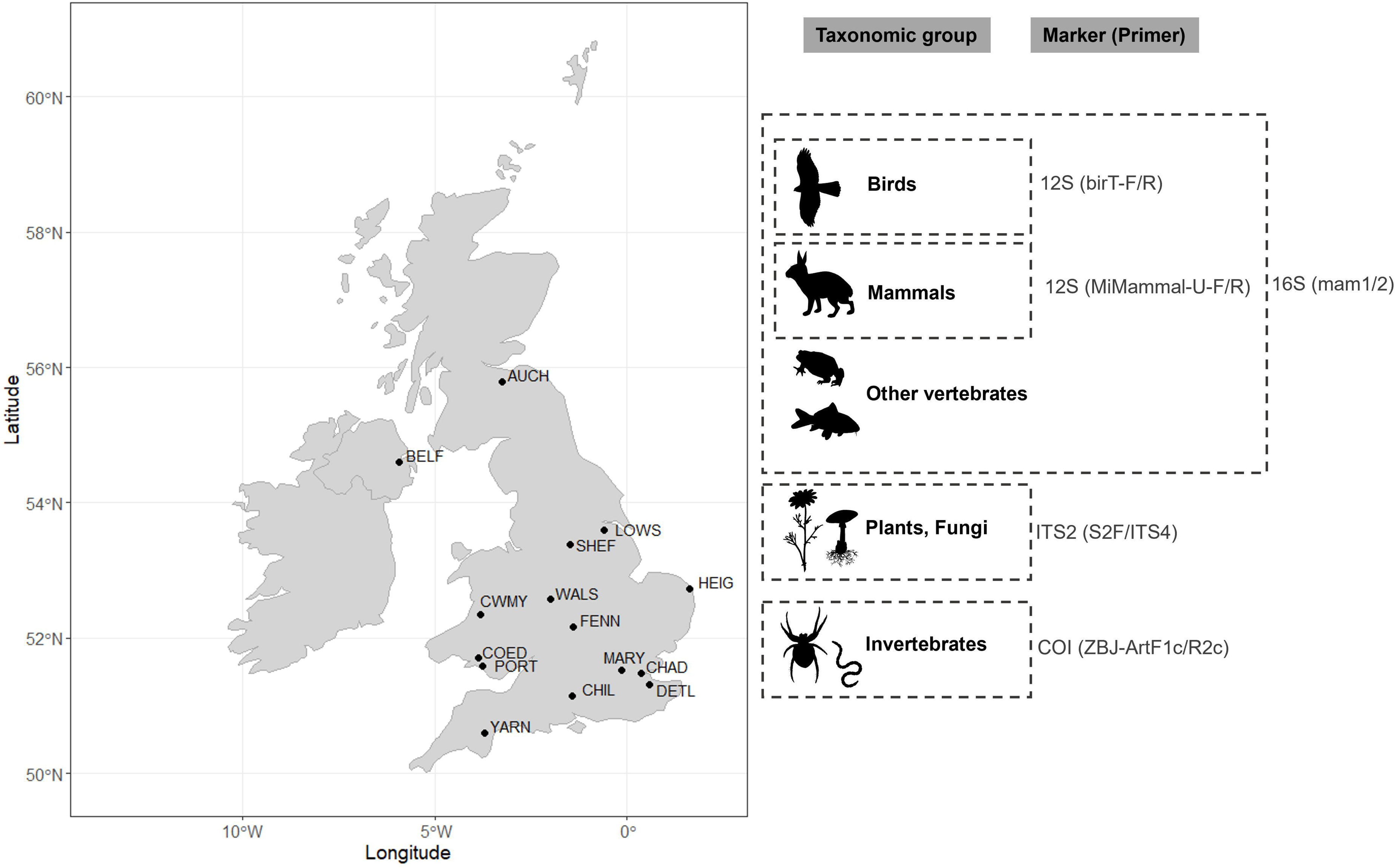
Map of the 15 sampling sites across the UK Heavy Metals monitoring network: one site in Northern Ireland (BELF = Belfast Centre), one site in Scotland (AUCH = Auchencorth Moss), ten sites in Great-Britain (LOWS = Scunthorpe Low Santon, SHEF = Sheffield Devonshire Green, WALS = Walsall Pleck Park, HEIG = Heigham Holmes, FENN = Fenny Compton, MARY = London Marylebone Road, CHAD = Chadwell St Mary, DETL = Detling, CHIL = Chilbolton Observatory, YARN = Yarner Wood), and three sites in Wales (CWMY = Cwmystwyth, COED = Swansea Coedgwilym, and PORT = Port Talbot Margam). The latitude and longitude of each site are publicly available on the UK AIR Air Information resource webpage (https://uk-air.defra.gov.uk/interactive-map?network=metals). The targeted taxonomic groups and associated markers and primer pairs are indicated on the right panel.

## RESULTS

### 1. A comprehensive terrestrial survey

Our dataset had a total of 18,373,005 reads which formed 1,556 ASVs that could be identified to unique taxa (details in Appendix S1 and Table S1). The ASVs represented a remarkable breadth of biodiversity, belonging to 1,220 genera of eukaryotes: 125 genera of vertebrates (1 phylum, 28 orders, 68 families, 136 species), 695 genera of invertebrates (7 phyla, 49 orders, 274 families), one genus of protist (3 phyla, 4 orders, one family), 210 genera of plants (6 phyla, 51 orders, 85 families) and 189 genera of fungi (4 phyla, 54 orders, 115 families) (Figure 2). The taxa recovered span a broad range of habitats, including terrestrial, sub-terrestrial (e.g. earthworms), semi-aquatic (e.g. waterfowls, water shrew, mayflies), as well as freshwater (e.g. common bream, stone loach), and marine (e.g. seabass, European hake) environments. We identified charismatic species such as newts, European robins, great tits, badgers and pipistrelle bats, and domestic and commensal species such as dogs, cats, donkeys, house mice and brown rats. We also identified invasive species such as Reeve’s muntjacs and Eastern grey squirrel, species of conservation concern such as the Eurasian skylark and European hedgehog, vectors of diseases such as mosquitoes, parasitoids and parasites such as fleas, ticks and pigeon lice. Invertebrate genera primarily belonged to insects and arachnids and included storage mites, leaf miners, gall inducers, potentially invasive and/or pest species such as *Harmonia* and *Drosophila,* and common nematodes of birds (*Ascaridia*). Finally, we recovered both native and ornamental trees (e.g. cypress, rhododendron) and plants (e.g. butterfly bush, geranium), crops (e.g. wheat, cucumber, brassicas), lichen and lichen-associated algae (e.g. *Trebouxia*, *Asterochloris*, *Chloroidium*), and pathogenic fungi (e.g. *Ramularia* on barley, *Leptosphaeria* on oilseed rape, *Hymenoscyphus* on ash). Two thirds of the vertebrates were diurnal (e.g. all birds but two, horse, squirrel), 12.5% were primarily nocturnal (e.g. tawny owl, bats), and 11.7% were both (e.g. cat) or crepuscular (e.g. deer, woodcock). Airborne eDNA captured the signal of widespread species (e.g. great spotted woodpecker, common pipistrelle) (Figure 3), as well as signals of species with a restricted distribution (e.g. bearded tit) or rarely visiting the UK (e.g. olive backed pipit) (Table S2). The most frequently detected taxa identified at the species or genus level were common plants (Figure S1A, > 50 detections: nettles *Urtica*, *Brassica*, ryegrass *Lolium*, meadow-grass *Poa*, wheat *Triticum* and soft-grass *Holcus*), common vertebrates (Figure S1B, > 50 detections: dogs, pigs, pigeons, magpies, pheasants, sheeps, robins, vole and badger), and diverse fungi (Figure S1C, > 50 detections; *Vichniacozyma* endophytic yeast, *Pichia* yeast, *Stemphyllum* causing common disease on UK crops, and *Leptospora*) and arthropods (Figure S1D, > 20 detections; *Psychoda* moth, *Cricotopus* and *Smittia* non-biting midges, psocopteran *Ectopsocus*, *Calliphora* and *Delia* flies, and *Entomobrya* springtails). Other invertebrates were less frequently detected (max. four detections; *Echiniscus* tardigrade and *Aulodrilus* annelid) (Figure S1E).

**Figure 2.**
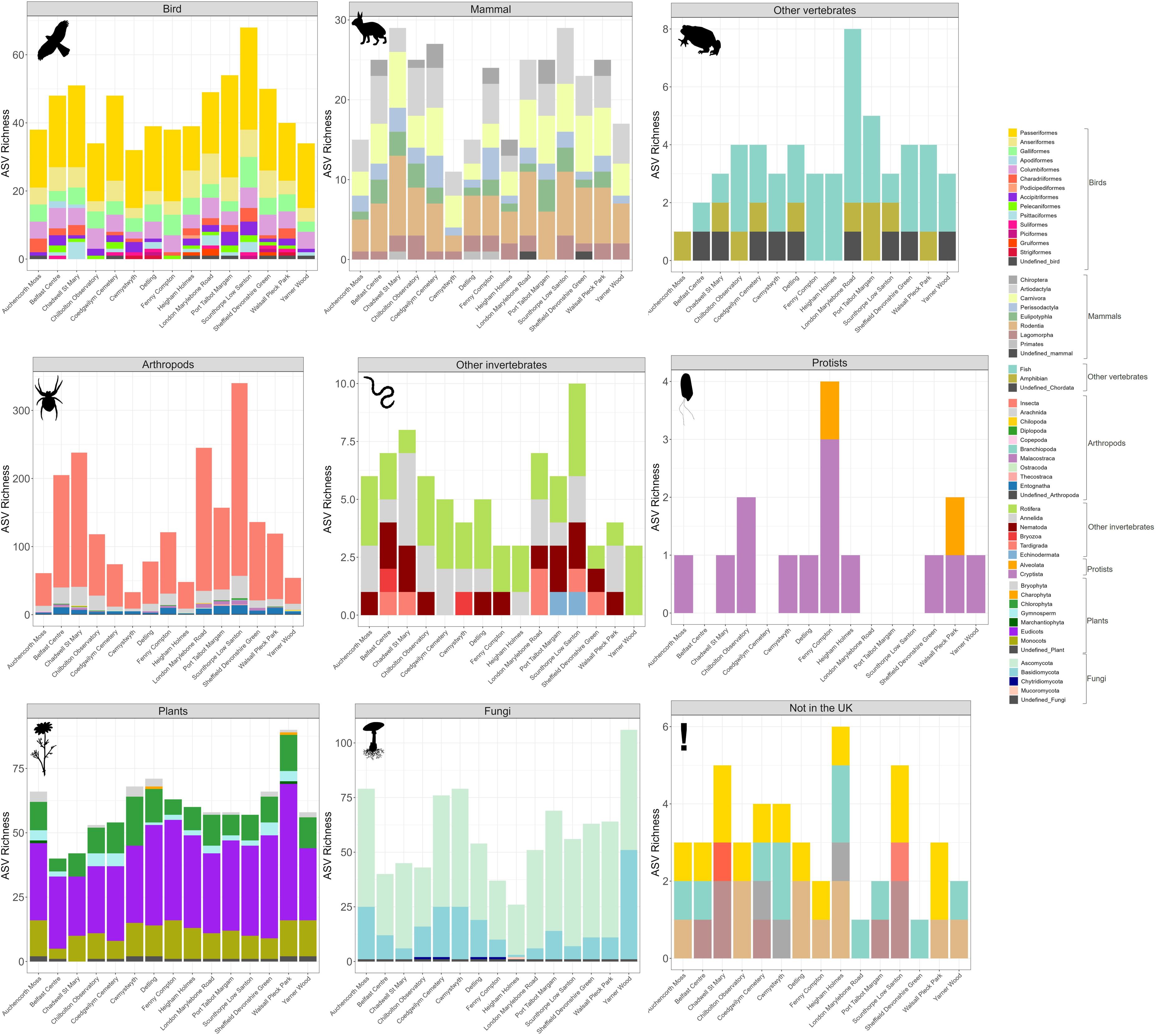
Representation of total ASV richness across sites and groups: Birds (Passeriformes, Anseriformes, Galliformes, Apodiformes, Columbiformes, Charadriiformes, Podicipediformes, Accipitriformes, Pelecaniformes, Psittaciformes, Suliformes, Piciformes, Gruiformes, Strigiformes, undefined), Mammals (Chiroptera, Artiodactyla, Carnivora, Perissodactyla, Eulipotyphla, Rodentia, Lagomorpha, Primates, undefined), Other vertebrates (fish, amphibian, undefined), Arthropods (Insecta, Arachnida, Chilopoda, Collembola, Diplopoda, Copepoda, Branchiopoda, Malacostraca, Ostracoda, Thecostraca, Entognatha, undefined), Other invertebrates ( Rotifera, Annelida, Nematoda, Bryozoa, Tardigrada, Echinodermata), Protists (Alveolata, Cryptista), Fungi (Ascomycota, Basidiomycota, Chytridiomycota, Mucoromycota, undefined), Plants (Anthophyta, Coniferophyta, Tracheophyte, Bryophyta, Chlorophyta, Marchantiophyta, undefined), and Not in the UK (bird, fish, mammal, Insecta). The 13 non-UK taxa (15 ASVs) likely originated from: 1) DNA contamination from the laboratory where the species are commonly handled (*Tachycineata bicolor*, *Trachops cirrhosus*, *Molossus, Melanoplus*), 2) lack of taxonomic resolution as similar UK species were also identified (*Alauda gulgula*, *Lepus californicus*, *Pipistrellus abramus*, *Garrulus lidthi*, *Chroicocephalus maculipennis*, *Barbatula to*ni, *Apodemus hermonensis*, *Acrocephalus orientalis*). The source origin of *Barbadocladius* is unclear.

**Figure 3.**
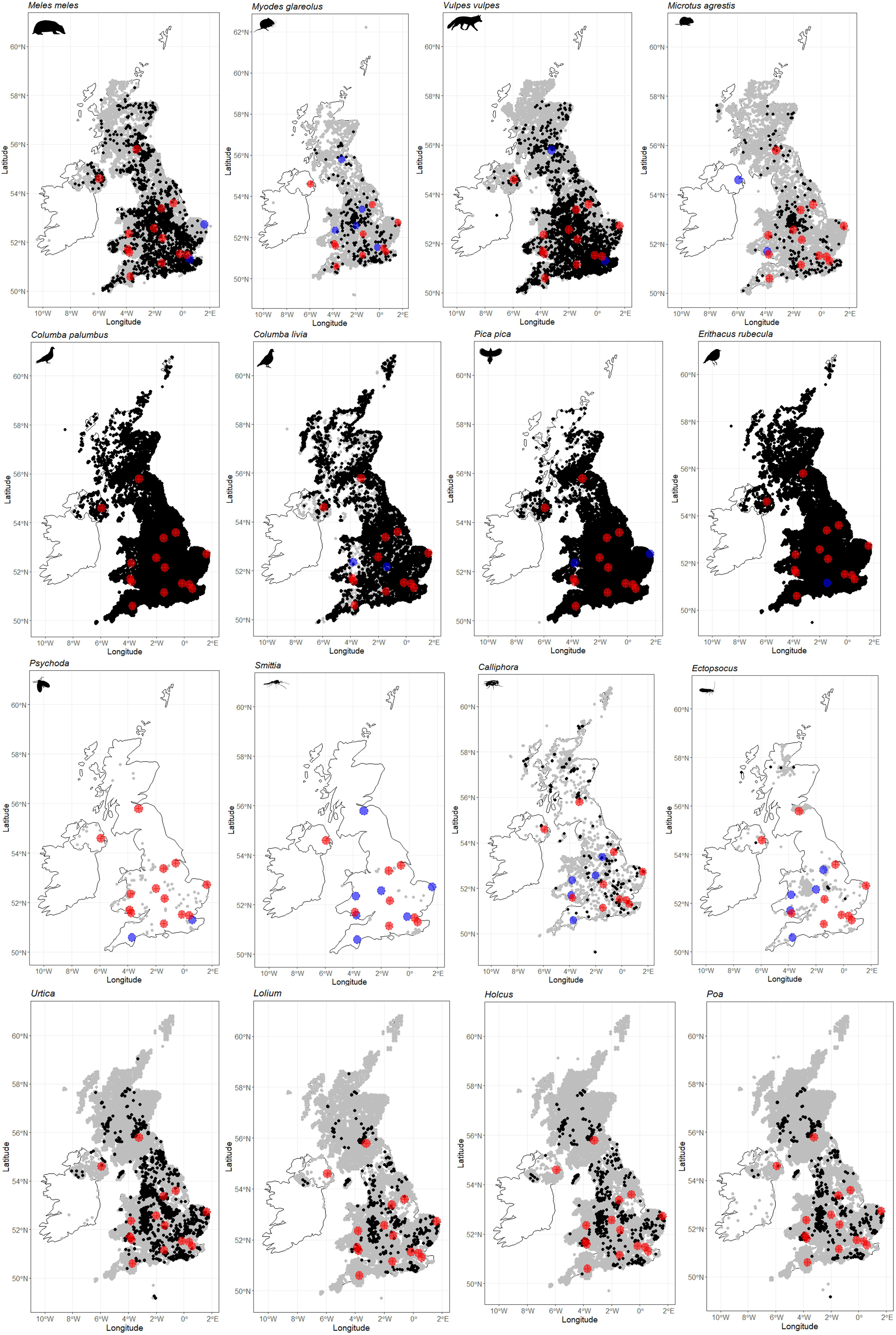
Map of the known distribution (NBN Atlas) of the top 4 most detected non domestic mammals, birds, arthropods and plants using airborne eDNA collected by the UK national ambient air quality monitoring network (185 eDNA samples at 15 sites from September 2021 to October 2022). The black and grey dots represent NBN Atlas confirmed observations for 2022 and all other years, respectively. Note that for arthropods, *Cricotopus* was in the top 4 but was not mapped because there were only unconfirmed records in NBN Atlas, and *Columba livia*, *Columba palumbus*, *Pica pica* and *Erithacus rubecula* had too many records in NBN Atlas (upper downloadable limit = 500,000 records), so only 2018 to 2022, 2022, 2021-2022, and 2022 records were used for these taxa respectively. The red crosses indicate the sites at which the taxa was detected using airborne eDNA, with a 18.6km buffer area shaded in red (i.e. median of the estimated transportation distances). The sites where eDNA was not detected are indicated in blue (18.6km buffer area). The maps were created using the ggplot2^78^, dplyr^80^, sf^83^ and rnaturalearth^84^ R packages. The NBN Atlas occurrence were downloaded at https://nbnatlas.org (accessed on 12 August 2024; data resource citations are available on FigShare using https://figshare.com/s/04229506308da822449c). All the PhyloPic silhouettes belong to the public domain, except for *Ectopsocus* (credit to Graham Montgomery, no changes, https://creativecommons.org/licenses/by/4.0/)

### 2. Several DNA markers are necessary to maximize biodiversity recovery

We observed a large overlap between the two markers (12S and 16S) used for amplifying vertebrate species, but each marker and primer pair recovered species not detected by the others (12S birds: 2, 12S mammals: 37 and 16S vertebrates: 33) (Figure S2). Overall, 12S mammals detected 74.3% of the vertebrate species, including mammals, birds, fish and amphibians. When combined with 16S vertebrates, the coverage increased to 98.5% and adding a third primer pair, 12S birds, resulted in a 100% coverage of vertebrate species within the dataset (Figure S2). The 12S mammals and 16S vertebrates both preferentially amplified mammal and bird species but also recovered amphibians and fish, while 12S birds only recovered birds. At the genus level, COI arthropods detected 57.5% of the taxa (arthropods, other invertebrates, vertebrates) within the entire dataset. Including ITS2 plants/fungi increased to 90.1% (plants, fungi, protists). Combining them with 12S mammals, we achieved 98% of genus recovery (Figure S2) within the entire dataset and adding 16S vertebrates to the three primer pairs allowed total recovery of the genera within the entire dataset. The 12S birds did not recover any additional genera beyond those detected by the other primers but identified unique species and showed good specificity for bird fauna which are frequent targets of biomonitoring programmes.

### 3. Airborne eDNA transportation

Among the 80 taxa whose presence in the UK was associated with human activities related to recreation (e.g. exotic pets, ornamental plants) and harvesting (e.g. agriculture) (see details in Table S2), we identified five exotic pets: the common parakeet, cockatiel, African grey parrot, peacock, and estrildid finches identified at the family and genus level (*Lonchura*). Their eDNA signals were found at distances ranging from 4.6 to 75.3 km to outdoor aviaries which could be potential sources of their eDNA in our samples. Two edible marine fish species, the seabass and European hake, were both detected in London at ∼400m from seafish stalls at the Marylebone Farmer’s market and 1.1km from the Church Street Market. Based on 16 comparisons (eight taxa) of detections to the closest known source (markets, aviary, etc; Table S2), distances ranged from 400m to 75.6km, with an average of the estimated maximum travel distances of 27.3km and a median of 18.6km. Finally, we identified four taxa not yet reported in England (invasive silver carp), not yet reported in Ireland (invasive crayfish *Pacifastacus*), or reported in the UK but with a lack of information on their distribution (arachnid *Nesticella*, invasive Prussian carp) (Table S2). We would classify these detections as tentative, given the lack of corroborating data to validate the record, but they suggest a reason for more intensive surveys in the local area of detection.

### 4. The complementarity of airborne eDNA and citizen-science databases

The eBird dataset identified more birds than airborne eDNA, with 257 species from 4,319 checklists compared to 83 species and six unique genera from the 185 eDNA samples. Across species, birds were detected at a higher number of sites using eBird than airborne eDNA (McNear’s Chi-squared *χ* = 511.9, p < 0.001), with 29.5% of the 1,392 species-site detections (i.e. unique species-site combinations) found in both the eDNA and eBird datasets (“Both”, Figure 4), 60.8% found only in the eBird dataset and 9.8% found only in the eDNA dataset. The most widespread birds (i.e. with the highest number of positive sites) were usually detected by both the eDNA and eBird approaches (e.g. robin *Erithacus rubecula*, common blackbird *Turdus merula*) (Figure 4). Some species were particularly well detected by eDNA (e.g. marsh tit *Poecile palustris*: 10 detections by eDNA only, one detection by eBird only, two by both), while others, such as the blue tit (*Cyanistes caeruleus*) and common kestrel (*Falco tinnunculus*) were never detected by eDNA (Figure 4). Similarly to eBird, the iNaturalist dataset comprised more taxa (1,421 taxa from 12,196 observations) than airborne eDNA (1,227 taxa from 185 samples). However, the overlap in shared species-site detections between the two datasets was lower (7.1%) than when comparing eBird and eDNA. Nonetheless, we observed higher overall detection rate by airborne eDNA than iNaturalist (McNear’s Chi-squared *χ* = 33.2, p < 0.001), with 43.0% found only in the iNaturalist dataset, and 49.9% found only in the eDNA dataset (Table S3). Airborne eDNA detected some taxa not recovered with iNaturalist (Branchiopoda, Copepoda, Ostracoda, Thecostraca, Nematoda, Rotifera, Tardigrada, Chytriomycota, Chlorophyta, Charophyta and Cryptista) (Figure 5, Table S3). Similarly, iNaturalist detected some taxa not recovered with eDNA (Reptilia, Mollusca, Cnidaria, Echinodermata, Platyhelminthes, Tunicata, Ceratophyllales, Eumagnoliids, Nymphaeales, Polypodiophyta, Chromista and Protozoa) (Figure 5, Table S3). While iNaturalist recovered more detections of Basidiomycota (*χ* = 18.8, p < 0.001), Bryophyta (*χ* = 7.2, p = 7.2e-03), Eudicots (*χ* = 232, p < 0.001) and Amphibian (*χ* = 7.7, p = 5.6e-03), airborne eDNA recovered more detections of Ascomycota (*χ* = 254, p < 0.001), Gymnosperms (*χ* = 25.7, p < 0.001), Annelida (*χ* = 8.4, p = 3.6-03), Entognatha (*χ* = 34, p < 0.001), Insecta (*χ* = 37.9, p < 0.001), Arachnida (*χ* = 6.5, p = 1.1e-02), bird (*χ* = 17.3, p < 0.001) and mammals (*χ*= 165.3, p < 0.001) (Figure 5). No significant difference was observed for fish (p = 0.371), monocots (p = 0.259), Bryozoa (p = 1.000), Malacostraca (p = 0.176), Diplopoda (p = 1.000), Chilopoda (p = 0.182) and Marchantiophyta (p = 0.130) (Figure 5).

**Figure 4.**
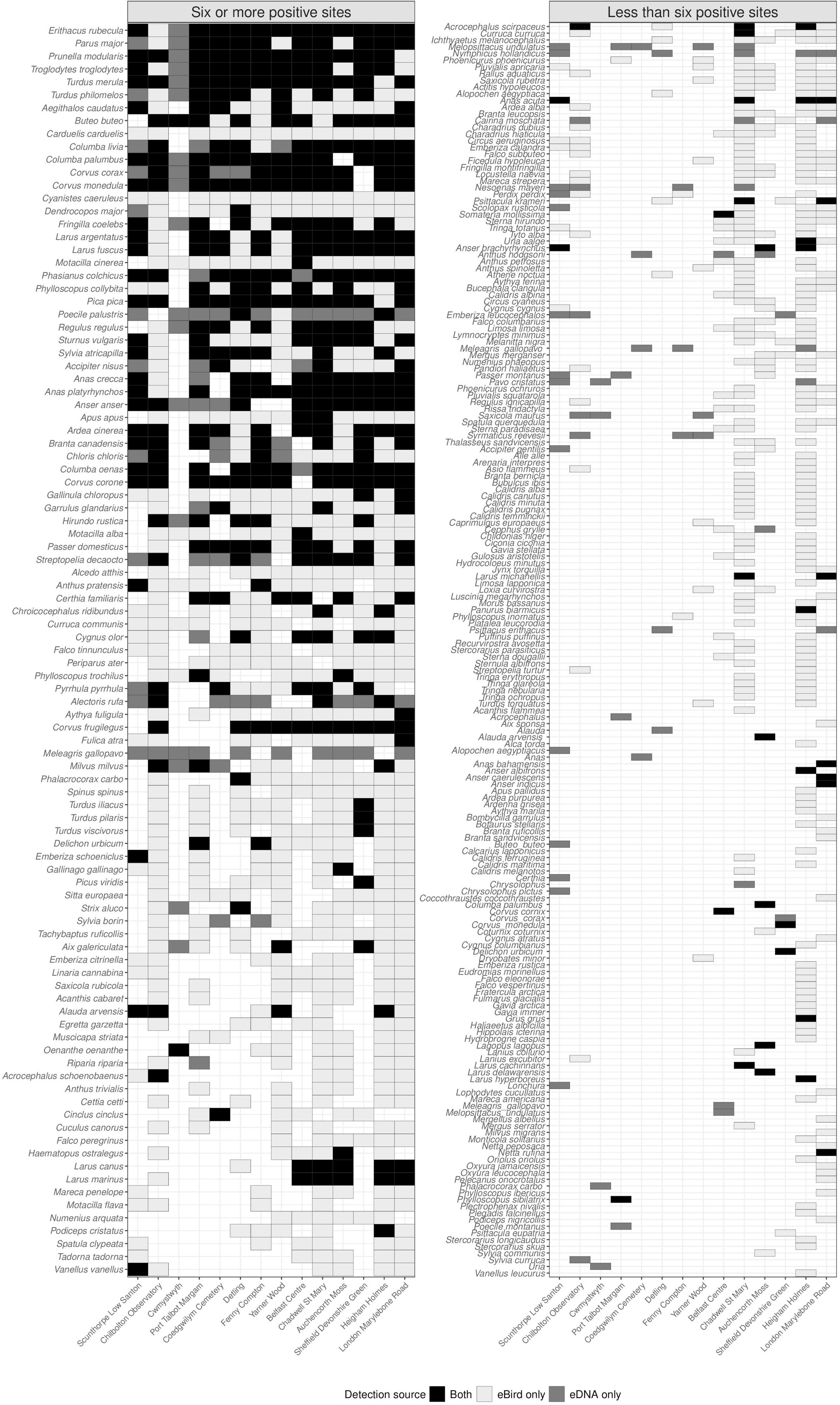
Bird detection source at each site, sorted from the highest to least number of detections (i.e. number of positive sites) from top to bottom. “Both” = detections in both the eBird and eDNA datasets, “eBird only” = detections in the eBird dataset only, and “eDNA only” = detections in the eDNA dataset only. eDNA data were obtained from a total of 185 eDNA samples, and eBird data from a total of 125 hotspots retrieved within a 5km radius of the sites: Scunthorpe Low Santon, Chilbolton Observatory, Cwmystwyth and Port Talbot Margam had 1 hotspot each, Coedgwilym Cemetery = 2,Detling = 3, Fenny Compton = 4, Yarner Wood = 8, Belfast Centre = 9, Chadwell St Mary = 10, Auchencorth Moss = 12, Sheffield Devonshire Green = 17, Heigham Holmes = 23 and London Marylebone Road = 33), including 4,319 checklists and 1,642 observers. Sites are sorted from the lowest to highest number of retrieved hotspots. Walsall Pleck Park was not included in the analysis as there was no eBird observation in a 5km radius of the site in 2022.

**Figure 5.**
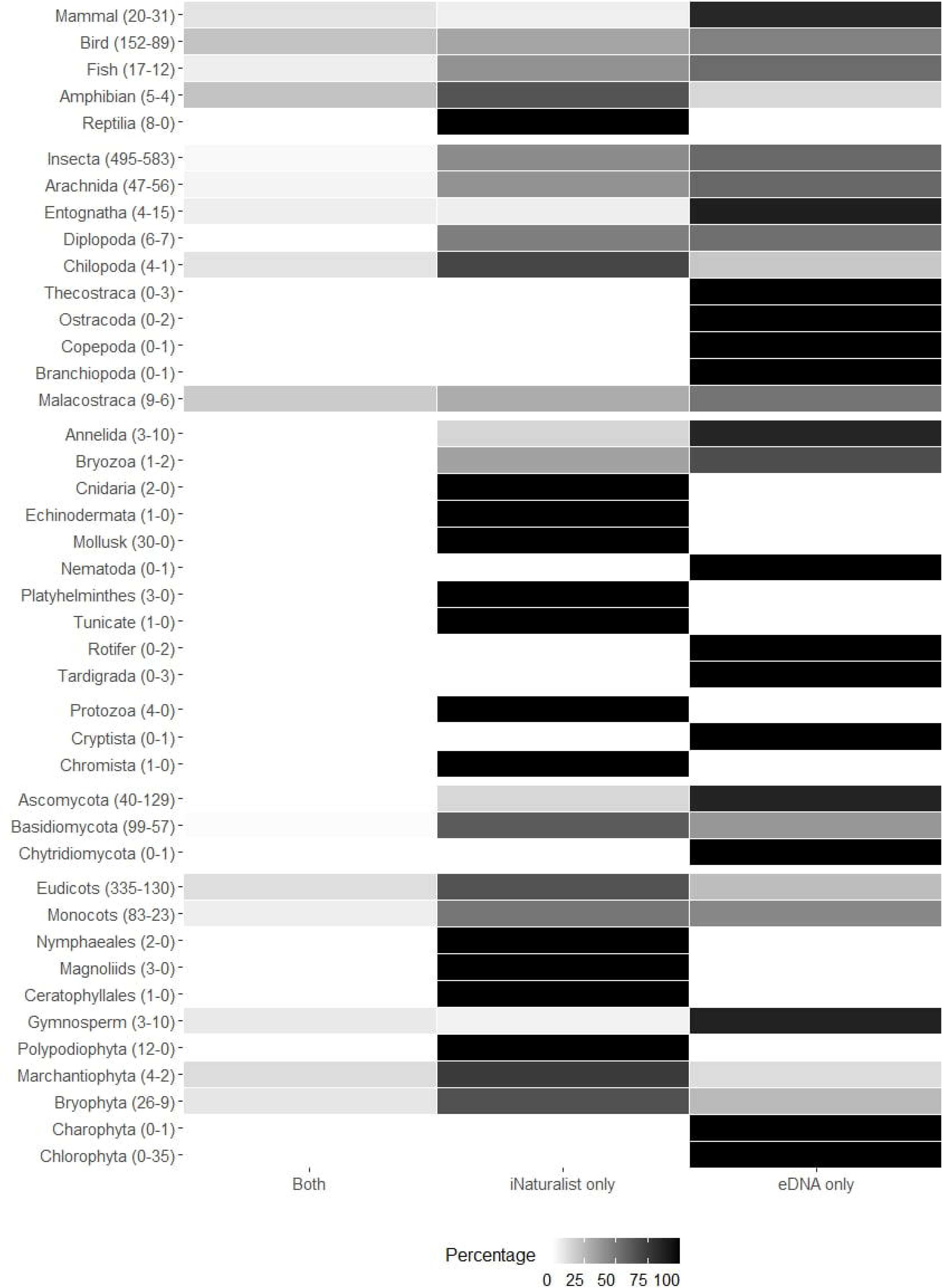
Detection source percentage per taxonomic group (vertebrates, arthropods, other invertebrates, fungi, plants and protists). “Both” = percentage of detections in both the iNaturalist and eDNA datasets, “iNaturalist only” = percentage of detections in the iNaturalist dataset only, and “eDNA only” = percentage of detections in the eDNA dataset only. The numbers of taxa in the iNaturalist checklists (total of 1,421 taxa from 12,196 observations) and the eDNA dataset (1,227 taxa from 185 samples) are indicated between brackets for each taxonomic group.

## DISCUSSION

Here we performed the first national survey using airborne eDNA and one of the first standardized, taxonomically general assessments of national terrestrial ecology. Using only pre-existing samples without any cold or chemical preservation that would optimize DNA recovery, and without any change to existing procedures in air quality monitoring, we were able to identify over 1,100 taxa at the species or genus level, spanning diverse taxonomic groups (vertebrates, invertebrates, protists, plants, fungi), body sizes (from a few micrometers to a meter), diel activity patterns, ecological niches and conservation statuses. While we treat all identifications with caution, given the dynamic nature of public reference databases of molecular data, and the small size of the fragments recovered, this broad spectrum of diversity confirmed that air pollution monitoring networks can efficiently capture terrestrial biodiversity, as well as some aquatic organisms^24,26,27^, to provide a realistic solution to monitoring the dynamics of life on land. The accuracy of such data will only increase as high quality global reference collections and related campaigns expand (e.g. Darwin Tree of Life www.darwintreeoflife.org, BIOSCAN https://ibol.org/bioscan/).

Airborne eDNA was also able to recover evidence from four vagrant bird species, difficult to observe in the field because of the rarity of their presence in the UK. Thus, airborne eDNA sampled by air quality networks could enhance the monitoring of common taxa but also elusive, historically understudied biodiversity and vagrant taxa and thus better align with global conservation goals^28^. Airborne eDNA could ultimately be applied beyond biodiversity monitoring as it successfully detected multiple confirmed invasive (e.g. Eastern gray squirrel), pest (e.g. weevils) and disease vector (e.g. mosquitoes, lice, ticks) species. We also recovered genetic evidence for three invasive species only very recently reported in the UK (Prussian carp in 2020), introduced in the UK but not yet reported in the area where it was detected with eDNA (*Pacifastacus* crayfish in Northern Ireland), or not yet reported in the UK (Silver carp). Such detections should be interpreted with caution as we anticipate errors in reference collections and often encounter IDs which are challenging to interpret at the species level given the small fragments of DNA being used. However, they suggest the potential of airborne eDNA to track invasion fronts, as previously demonstrated with aquatic eDNA^29,30^, and provide a reasonable line of evidence for enhanced surveys in those locations to confirm or refute our observations.

The eDNA collected on the filters likely originated from both organismal (e.g. microalgae, unicellular fungi) and extra-organismal bioaerosols^31^, including excretions (e.g. brochosomes produced by insects, faeces) and fragments from plant debris, hyphae, hairs, feathers, eggs or scales, bound or not to ambient particles^32–34^. It is possible that eDNA also originated from propagules released into the air by fungi and plants but as PM_10_ have a diameter equal or less than 10 µm, many pollen grains and spores would be excluded. National sampling networks use different size selective sampling heads to collect either PM_2.5_ or PM_10_ (particles ≤ 2.5 and ≤ 10 µm, respectively) which may both capture eDNA but with different properties. Our choice of PM_10_ was deliberate, beyond the advantages of having a higher number of PM_10_ samplers within global ambient air quality networks. Specifically, targeting smaller size fractions (e.g. PM_2.5_) could improve the recovery of total DNA by capturing more subcellular material^35–37^ but may penalize some taxonomic groups such as plants and fungi. More importantly, the smaller the particle size, the greater the travelling distance in the air, which would limit the spatial resolution and the interpretation of eDNA signals in biodiversity surveys. For example, so-called “Sahara dust events”, transporting dust intercontinentally, specifically occur at the lower size fraction. PM_10_ is therefore a good trade-off as it likely recovers subcellular and cellular materials while limiting transboundary transportation. Further work evaluating the influence of particle size and sampling head on taxa recovery and transportation distance is needed to optimize airborne eDNA approach as transport distance will have implications for the scale of observation that can be recovered for conservation purposes. While wind transportation is a strong argument in favour of large particle sizes for analysis^22,24,38,39^, the structure of the particles and therefore their aerodynamics properties can also have an influence^40^ and should be further investigated.

Our estimations of airborne eDNA transportation distance ranged from a few hundred meters to 75.3km, supporting previous suggestions of limited airborne eDNA transportation from relatively local sources (≤ 100km)^25,41^. Transportation distance (estimated median distance of 18.6km) is likely to be even less as it was difficult to determine all the possible sources of eDNA from internet searches (e.g. there could be private aviaries that could contribute DNA but are not publicly available on maps) and the analysis was based on a limited number of taxa. Moreover, as the air quality monitoring networks were established to monitor health effects of air pollution, the samplers are near ground in the typical human breathing zone of ∼1.5-2m. Thus, eDNA travelling distance is likely to be significantly constrained by local topography and landscape^42,43^, with long transport possible but not predominant, providing a veritable “sweet spot” for mapping biodiversity. Estimates will be refined as more data are analysed but already provides a promising range and scale for national level biomonitoring.

Previous airborne studies showed contrasted results about the effect of sampling time, sampling duration and weather conditions on eDNA detectability in the air^24,26,31^. Using air pollution monitoring infrastructure, the air was sampled day and night almost continuously for a year, increasing the probability of detecting organisms regardless of potential eDNA diurnal pattern and meteorological conditions (e.g. rain) thereby increasing the taxonomic scope of our detections. The detection of a few aquatic organisms was also not unexpected as previous airborne eDNA studies recovered fish and amphibian DNA too^24,26,44,45^. The capture of aquatic species, such as zooplankton, in the air has also been observed with traditional methods^46,47^. While we cannot determine the exact mechanisms underlying these eDNA detections in our dataset without a targeted experimental design, several non-exclusive plausible scenarios could explain their presence. For example, windy conditions can resuspend top-soil and water surface particles, facilitating the detection of ground-dwelling, semi-aquatic, aquatic and subterrestrial organisms such as the earthworms^25^. Like rivers, wind could thus be considered as a conveyor belt of biodiversity information^48^. Secondary sources, such as diet remains (e.g., prey DNA or fragments on bird feathers or mammal fur), could also carry trace amounts of aquatic DNA into the air. Finally, our results showed a long persistence of taxonomically diverse eDNA on the filters despite a storage at ambient temperatures (standards of heavy metal analyses for which they originally collected) for up to 22 months before being moved to -20°C. It is likely that the dry and light-proof storage conditions (petri dishes in light-proof cupboards) until freezing reduced eDNA degradation effects associated with UV exposure, microbial activity and contact with ambient air.

Although our samples were not appropriate for temporal analysis of biodiversity (see Appendix S2), better sample preservation (e.g. immediate freezing) will enable users to exploit the full data potential associated with the high temporal replication (e.g. seasonality) produced by air pollution monitoring infrastructure.

Environmental DNA surveys often outperform traditional surveys^49–51^ but we found complementary results among taxonomic detections recovered by airborne eDNA and citizen science databases. While both dataset types are biased in their own way (e.g. variations in sampling effort), they are an important comparison because citizen science provides one of the rare datasets with comparable temporal and spatial coverage. The performance of only 185 airborne eDNA samples is extremely promising despite suboptimal storage conditions considering the sampling effort of the two citizen databases (thousands to tens of thousands of observations). As expected, the generalist iNaturalist citizen-science program recovered numerous charismatic taxonomic groups relatively easy to spot and/or identify morphologically (e.g. amphibians, reptiles, snails, club-shaped fruiting-bodied fungi, plants)^52–54^. Both citizen science databases had a higher number of bird species than airborne eDNA, likely due to the emphasis on birding in both UK wide campaigns (e.g. Big Garden Birdwatch count) and the general popularity of birding. Within a 5km radius around the sites, iNaturalist did not recover more detections than airborne eDNA despite higher overall taxa richness, likely due to its opportunistic and generalist nature: while it detects a greater diversity of taxa, these detections are not always consistently recorded within close proximity to the sites, especially on a large scale (e.g. birds, Figure 5). By the nature of its development as a national monitoring infrastructure, airborne eDNA was overall more efficient in documenting a broader spectrum of diversity (including communities more numerous on average at night such as insects^55^) and at mapping some taxa over large scales with limited sampling effort.

However, as the radius around the sites increased to 18.6 km (the tentative estimated distance eDNA can travel here), iNaturalist showed higher detection rates for most tested taxonomic groups (see Appendix S3). This highlights the effectiveness of massive citizen science efforts in gathering broad-scale biodiversity data and underscores the need for more accurate estimations of eDNA transportation distances to further improve comparisons between biodiversity monitoring approaches.

While quantifying abundance or biomass from eDNA is challenging^56^, it is reassuring that the most frequently detected species in our dataset commonly occur in the UK (National Biodiversity Network Atlas records) and our detections matched the known presence of species at the sites (Figure 3). For example, the Auchencorth Moss sampler recovered dog DNA likely from a dog park 1.1km away and walked dogs, horse DNA likely from the horse farm 400m away, and wild species DNA found at the Leadburn Community woodland 2.2km away (e.g European hare, greylag geese, common toad, willow, cuckoo flower, clover, foxgloves). Exploring the data being excluded by filtering with the negative controls and ghost plates revealed the occurrence of some organisms seen occasionally at the site (e.g. red fox, European rabbit, sparrowhawks, common pipistrelle). It is preferable to remove such low-level background noise, despite detections making sense, to avoid ascertainment biases and false positives. However, this suggests the value of learning about the analytical processes, for example by exploring the effect of using multiple thresholds^57^. Indeed, filtering metabarcoding data is absolutely necessary to remove false positives and standardize data processing for improved reproducibility^58^, but it also inflates the rate of false negative detections and should be interpreted cautiously^23,59^. Yet, the majority of false negative detections were not associated with ASV filtering, with some taxonomic groups never detected by our eDNA approach (e.g. reptile, mollusk, odonate, fern). This might be explained by 1) their low DNA shedding rate limiting the probability of eDNA capture on the filter (rarity of template DNA), 2) the size selective sampling head preventing large particles to reach the filter (e.g. fern spore), 3) incompleteness of reference databases limiting taxonomic assignment of ASVs, 4) primer choice, as for example our vertebrate primer pairs were initially designed for mammals and birds and have therefore less affinity to other vertebrates, or 5) competition between various DNA templates during PCR^58,60^. This strongly supports the use of multimarker approaches to maximize taxonomic coverage in airborne eDNA surveys^22,24^. The presence of false negatives also suggests that sampling and processing strategies could be optimized for a maximized airborne eDNA capture and identification, and that combined approaches (e.g. metabarcoding + shotgun sequencing^61^) are likely optimal to maximize coverage. Air pollution monitoring samplers collected one filter per week, but in this case four quarters (one month) had been stored together, with each quarter of the filter representing a week of continuous air filtering. The quarters were thus considered as one “sample” with cross-sharing of taxa by physical contact of the filters, but pooling and extracting the four quarters separately would maximize the capture of eDNA diversity^62^. We also recovered 15 ASVs not occurring in the UK whose presence in our dataset could originate from 1) DNA contamination from the laboratory where the species are commonly handled (e.g. *Tachycineata bicolor*) or incorrect identification by our markers as we also detected closely related species (e.g. *Corvus dauuricus*). The non-UK taxa represented only 1.0%, 1.1% and 0.9% of the total number of ASVs, reads and detections, respectively. Nonetheless their presence highlights the importance of carefully checking the lists of taxa as this can reveal false positives from several origins. We used a manual curation approach checking ASV identifications one at a time, though this approach does not scale. For our largest datasets (plants/fungi and arthropods), we performed a classification followed by a manual checking for subsets to try and streamline the approach. In these cases, we also limited the identifications to genus to avoid inflated false positives from auto classifications. While not ideal, and in many cases species level ID was likely possible, this conservative approach provides an alternative solution to handle large datasets in a reasonable amount of time while minimizing errors in taxonomic assignment.

In conclusion, airborne eDNA collected from existing semi-automated air monitoring infrastructures offers a comprehensive, cost-efficient method for surveying terrestrial biodiversity and fulfilling the goals of the Strategic Plan for Biodiversity and Sustainable Development Goals. This scalable solution to biodiversity assessment relies on international protocols which have already been highly standardized (e.g. sampler head and sampler height, volume of air filtered, filter type), favoring consistency and comparability across studies. Further methodological studies will be particularly useful to improve our understanding of airborne eDNA transportation, degradation and preservation, as well as the effect of particle size, shape and origin on the recovery and identification success of the taxa. Similar to aquatic eDNA studies for aquatic biodiversity, comparing airborne eDNA and traditional surveying methods (e.g. malaise traps) will help assess detection efficiency, taxonomic coverage, and potential biases, ultimately strengthening its role in biodiversity monitoring. While we successfully captured a swath of diversity across the UK, including under-represented taxa in traditional biodiversity surveys, optimizing filter storage and processing will improve detection and taxonomic coverage. Additionally, the detection of invasive species, pest and diseases vectors highlights the potential applications beyond biodiversity monitoring. As a powerful tool for assessing biodiversity baselines and changes, airborne eDNA collected by air monitoring infrastructures could support regional and national decisions and eventually enable the exploration of the three levels of biological diversity: species, ecosystems and genetics^18,63^. In the face of the current global biodiversity crisis, expanding and integrating diverse monitoring approaches, particularly making use of existing large scale infrastructures, will be essential to achieving conservation goals.

## METHODS

### 1. Sample collection

Airborne eDNA was collected between September 2021 and October 2022 using 15 Digitel DPA-14 ambient air samplers on the UK Heavy Metals monitoring network^39^ (Figure 1). The latitude and longitude of each site are publicly available on the UK AIR Air Information resource webpage (https://uk-air.defra.gov.uk/interactive-map?network=metals). The semi-automated samplers of PM_10_ fraction sampled air at a rate of 1 m^3^/h for four periods of seven consecutive days over a 28-day period. During each seven-day period, one 47 mm diameter cellulose ester filter (Merck MF-Millipore, 0.8 µm mixed cellulose esters membrane) was sampled following the procedure developed for the UK heavy metals monitoring network.

After each week, the new filter was translated into position. After the collection period, the filters were returned to the National Physical Laboratory (NPL) where they were kept refrigerated for an average of two weeks before a quarter of each filter was sub-sampled (cut using a pair of clean ceramic scissors against a circular cutting template marked with four quadrants), and analysed for heavy metals. The remaining filters were stored in Petri dishes (samples separated by location and month of sampling) and kept in light-proof cupboards in ambient conditions up to 22 months before quarters were subsampled, transferred to zip-lock bags and stored at -20°C at McGill University (Canada) in July 2023. We also included one unused filter per site (except London Marylebone Road and Auchencorth Moss) as field controls. This control filter was placed in the bodywork of the sampler but was not used for actual air sampling^26^.

### 2. DNA extraction, library preparation and sequencing

DNA extraction and PCR mix preparation were conducted in a dedicated pre-PCR laboratory, with separate rooms assigned to each processing step (details in Appendix S1). One quarter of each filter (N =185 eDNA + 13 control filters) was transferred to sterile 2 mL tubes using sterile tweezers. DNA was extracted using the Qiagen Blood and Tissue kit following the manufacturer’s protocol with slight modifications described in Appendix S1.

Negative controls of extraction (only reagents) were included approximately for every 11 samples (N = 19). We also included three filters that were not used or placed in the sampler but were simply stored in zip-lock bags as filter blanks.

We amplified five regions targeting vertebrates (mitochondrial 16S), mammals (mitochondrial 12S), birds (mitochondrial 12S), arthropods (mitochondrial COI), and plants and fungi (nuclear ITS2), using the mam1/2 primers^64,65^, MiMammal-U-F/R primers^66^, birT-F/R primers^67^, ZBJ-ArtF1c/R2c primers^68^ and S2F/ITS4 primers^69^, respectively (Figure 1, Appendix S1) and included six PCR negative controls. PCR_1_ (including samples, negative and positive controls) were run in three technical replicates. Detailed protocols of library preparation are available in Appendix S1. All the samples and controls were sequenced on Illumina NextSeq2000 sequencer (2x300 PE cycle run, P1 reagent kit) at the Canadian Centre for DNA Barcoding (CCDB, Guelph, Ontario) (Appendix S1).

### 3. Processing of the sequencing data

All steps and used parameters are described in detail in Appendix S1. Briefly, R1 and R2 reads were assembled using the PEAR v0.9.11 program^70^ and Cutadapt v3.7^71^ was used to demultiplex assembled reads by sample and remove M13 adapters (CCDB). The full index combinations were searched during the demultiplexing step even if they were not used in our project (“ghost plates”) to estimate sample assignment errors. The merged reads were split by primer pair with Cutadapt 4.5 and processed using the DADA2 pipeline v1.26.0^72^ in R 4.4.0^73^. Reads were truncated after 157 bp (COI arthropods), 171 bp (12S mammals), 90 bp (16S vertebrates), 265 bp (12S birds) or 230 bp (ITS2 plants/fungi), and reads matching against the phiX genome, with ambiguous bases, or with more than two expected errors were discarded. All identical sequencing reads were then combined into unique sequences (dereplication), ASVs were made using the learnt error rate and chimeras were removed.

Taxonomic assignment was kept at the species level only with query cover > 60% and percentage of identity ≥ 97%, and at the order level with percentage of identity < 97% (Appendix S1). In cases where multiple species matched with the same percentage of identity, the closest common ancestor (e.g., genus) was retained. For the 16S and two 12S markers we used the online BLAST tool (NCBI full nucleotide collection) to manually assign taxonomy to the ASVs. For the two markers with large numbers of ASVs (4,389 and 7,550 ASVs for ITS2 and COI, respectively) we combined automated non-Bayesian taxonomy classifier (sintax command of the usearch v11.0.667 pipeline^74^) and manual (online BLAST tool, NCBI full nucleotide collection) approaches (Appendix S1). We used sintax against the BOLD database for COI, and against two databases PLANiTS_29_03_2023^75^ and Unite all eukaryotes v10.0^76^ for ITS2; with a probability threshold of 0.60 (see details on the taxonomic assignment strategy in Appendix S1). Because invertebrates, plants, and fungi are less documented (known occurrences and reference databases), and the species lists are harder to validate due to short fragment size, the large number of ASVs and lack of specialized taxonomic expertise in these groups, only genus-level identifications were retained for ITS2 and COI markers^26^.

### 4. Filtering assigned ASVs

The 185 samples were merged by site (N=15) because of the negative effect of ambient storage time on ASV recovery (see Appendix S2). Non-identified ASVs (“no match”) and ASVs identified as common suspected contaminants before any filtering of data were removed^23,26^. For the vertebrate and invertebrate regions, discarded ASVs included human, bacteria, Amoeba, *Gallus gallus* (chicken), *Bos taurus* (cow) and non-animals (plants, rhodophytes, oomycetes, heterokonts, fungi). For the plants and fungi region, we removed Metazoa, *Alternaria*, and *Penicillium*. Retained ASVs were filtered using the reads wrongly assigned to non-used index combinations (‘ghost plates’): we kept an ASV detection only if its read count was greater or equal to its highest read count in the ghost plates. Finally, ASVs detections with a read count below their highest read count in the negative controls were excluded. ASVs with identical identification were merged (sum of the reads). Then, vertebrate data from the 16S and two 12S markers were merged. Finally, taxonomic assignments were classified as follow: “Native or established’’ for species present in the UK, “Visitor’’ for rare visitors from the continent, “Human-associated” for species not naturally occurring in the UK but that can be found due to human activities (e.g. recreation, farming), “Insufficient information” for species lacking information on their presence in the UK, and “Not in the UK” for species not occurring in the country (e.g. wrong molecular identification due to marker resolution).

### 5. Analysis

We investigated the impact of marker choice on vertebrate detections (species level; the 16S and two 12S markers), and on all taxa detections (genus level; all markers) by generating Venn diagrams and assessing cumulative primer pair coverage using the online SNIPe tool^77^. We visualized ASV richness across sites and groups (birds, mammals, other vertebrates, arthropods, other invertebrates, protists, plants, fungi, non-UK taxa) in R4.4.0 and Rstudio 2024.04.1+748 using the following packages: ggplot2^78^, tidyr^79^, dplyr^80^ and stringr^81^. Then, the taxa in the “Not in the UK” category were excluded from further analysis. The same R packages were used to investigate the number of detections (positive samples) of each vertebrate, arthropod, other invertebrate, fungi and plant taxon. We used six taxa in the “Human-associated” category and two native species (Soprano pipistrelle and Whiskered bat) to estimate airborne eDNA transportation distances by measuring the distance between the sites at which they were detected and their closest known location (“Measure distance and areas” tool in Google My Maps). Because the true source origin(s) of these species’ DNA could be missed, the estimated distances should be interpreted with caution. This analysis provides a preliminary assessment, but a dedicated experimental design will be necessary to further rigorously test eDNA transport distances. Humans-associated plants were not included in distance calculations as their widespread distribution makes it difficult to determine a single source. The approximate median estimated transportation distance was used to map a buffer area around the sites with positive eDNA detections of the top four most detected mammals, birds, arthropods and plants on their known distribution. The known distributions were mapped using NBN Atlas confirmed records for 2022 and all available other years^82^. Note that the arthropod *Cricotopus* was in the top four but was not mapped because there were only unconfirmed records in NBN Atlas, and *Columba livia*, *Columba palumbus*, *Pica pica* and *Erithacus rubecula* had too many records (upper downloadable limit = 500,000 records), so only 2018 to 2022, 2022, 2021-2022, and 2022 records were mapped for these taxa, respectively. Downloaded NBN Atlas records and data resource citations are available on FigShare using https://figshare.com/s/04229506308da822449c. The maps were created using the ggplot2^78^, dplyr^80^, sf^83^ and rnaturalearth^84^ R packages.

Finally, we compared our lists of taxa obtained with airborne eDNA at each site to publicly available records in citizen science databases as they are currently among the only datasets with such comparable temporal and spatial coverage. We selected two citizen science databases: eBird (www.ebird.org) specializing in birds and iNaturalist (https://www.inaturalist.org/) covering animals, plants and fungi. We included taxa belonging to all categories but “Not in the UK”. In eBird, we used the “Explore Hotspots - eBird” map to retrieve observations from January 1^st^ to December 31^th^, 2022 within a 5km radius around each sampler (maximum of 33 hotspots per site), resulting in 4,319 checklists from 1,642 observers at 125 observation hotspots). In iNaturalist, we used the “Export observations’’ tool with the following parameters: period: January 1^st^ to December 31^th^, 2022, Longitude/latitude coordinates of the air sampler with a 5km radius, “Research” quality grade, any taxa but unknown, “Yes’’ to verifiable, and “Any” to “Threatened”, “Introduced”, “Native” and “Popular” (1,421 taxa from 12,196 observations). For each analysis we used three categories: “eBird/iNaturalist only” for taxa only identified in the citizen science species list, “eDNA only” for those only identified in the eDNA lists, and “Both’’ for those identified in both the citizen science and eDNA datasets. We included ASVs identified at the genus level when: i) zero or one species belonging to the genus was detected in the citizen science dataset at the site (e.g. *Uria* and *Passer* at Cwmystwyth), ii) several species were identified in the citizen science dataset and ASVs were identified at both the species and genus level (e.g. *Anas*) or genus only (e.g. *Larus*). We used the mcnemar.test function to perform a McNemar’s Chi-squared test comparing detection rates of eBird vs eDNA and iNaturalist vs eDNA methods. Since iNaturalist and airborne eDNA detect a broad range of taxonomic groups, we also conducted this test for each taxonomic group where a suitable contingency table was available (i.e., with at least one discordant pair indicating detection by one method but not the other). The results of the analysis for iNaturalist detections at an 18.6 km radius around the sites are presented in Appendix S3. Note that we did not conduct the analysis for eBird detections at the 18.6 km radius, as it was already outperforming eDNA at the 5 km radius.

## ACKNOWLEDGMENTS

We are grateful to Chris Robins, Vandana Kantilal, Emma Braysher and Jody Cheong for collecting and preparing the filters, Daniel Marquina for helping with DNA extractions, Evgeny V. Zakharov for managing the sequencing, and Robin Floyd for helping with taxonomic assignments. This work was supported by The Natural Sciences and Engineering Research Council of Canada through the Discovery Grants Program, The Government of Canada’s New Frontiers in Research Fund (NFRFT-2020-0073) Tracing the Patterns of Life on a Changing Planet, Genome Canada (BIOSCAN Canada) and Ontario Genomics (OGI-208), Future Leaders Fellowship (MR/Y016971/1) and Canada Graduate Scholarships - Doctoral (CGS D) program. Collection and processing of the samples was funded by the Environment Agency, the UK Department for Environment, Food and Rural Affairs, and the UK Department for Science, Innovation and Technology.

## AUTHOR CONTRIBUTIONS

Design of the study: E.L.C, O.T., M.E.C, J.E.L, J.J.A, A.S.B

Sample collection: J.J.A, A.S.B Labwork: O.T.

Bioinformatics: O.T., N.R.G, E.L.C Analyses: O.T.

Drafting of the manuscript: O.T. Editing of the manuscript: all authors

Funding acquisition: J.J.A, A.S.B, M.E.C, E.L.C

## DATA AVAILABILITY STATEMENT

All sequence data (reads demultiplexed by sample and primer pair) have been deposited in NCBI Sequence Read Archive (PRJNA1148205) and are publicly available as of the date of publication. Downloaded NBN Atlas records and data resource citations are available on FigShare using https://figshare.com/s/04229506308da822449c.

