## Appendix S1 for "First national survey of terrestrial biodiversity using airborne eDNA"

**Appendix S1.** Detailed laboratory and bioinformatic protocols, and sequencing results

*Laboratory conditions*

DNA extraction and PCR preparation were conducted in a dedicated pre-PCR laboratory, with separate rooms assigned to each processing step. Personnel working in the pre-PCR lab wore coveralls, masks, hairnets, and disposable nitrile gloves to minimize contamination. PCR amplification took place in a designated PCR/post-PCR laboratory.

To prevent contamination, all bench spaces and laboratory equipment were thoroughly cleaned with a 20% bleach solution, followed by rinsing with distilled water. Aerosol barrier filter tips were consistently used to prevent cross-contamination between samples.

*Modifications of the DNA extraction protocol*

We followed the Qiagen Blood and Tissue kit manufacturer’s protocol for tissue with the following modifications: 1) we added 540 µL of ATL buffer and 60 µL of proteinase K to the tubes and shook them at 160 rpm during overnight incubation at 56°C, 2) we used 600 µL of buffer AL and 600 µL of ethanol, and 3) we added 60 µL of AE buffer directly onto the pad, centrifuged after a 5-minutes incubation, and pipetted the 60 µL eluate onto the pad again for another 5 minutes incubation before final centrifugation.

*Detailed library preparation*

PCR1 were performed in 15 µL reaction volumes using 7.5 µL of Qiagen Master Mix (2X), 0.25µL (10µM) of each primer, 4 µL of UltraPure DNase/RNase-Free Distilled Water (ThermoFisherScientific) and 3 µL of DNA. We amplified five regions targeting vertebrates, mammals, birds, arthropods, and plants and fungi. For vertebrates, we used the mam1/2 primers^1,2^ (mam1: 5’-CGGTTGGGGTGACCTCGGA-3’; mam2: 5’-GCTGTTATCCCTAGGGTAACT-3’) to amplify a ~90bp region of the 16S gene. The thermocycling conditions were: 95°C for 15min, followed by 40 cycles of 94°C for 30secs, 55°C for 90secs, 72°C for 90secs, and a final extension of 72°C for 10 mins. To target mammals, we used the MiMammal-U-F/R primers^3^ (MiMammal-U-F: 5’- GGGTTGGTAAATTTCGTGCCAGC-3’; MiMammal-U-R: 5’-CATAGTGGGGTATCTAATCCCAGTTTG-3’) amplifying a 171bp region of the 12S gene. The thermocycling conditions were: 94°C for 5 min, five cycles of 94°C for 1 min, 45°C for 1 min 50 sec, 72°C for 1 min, followed by 35 cycles of 94°C for 1 min, 60°C for 1 min 50 sec, 72°C for 1 min and a final extension at 72°C for 10min. For birds, we amplified a ~277bp region of the COI gene with the birT-F/R primers^4^ (birT-F: 5’-YGGTAAATCYTGTGCCAGC-3’; birT-R: 5’-AAGTCCTTAGAGTTTYAAGCGTT-3’). The thermocycling conditions were: 95°C for 10min, followed by 35 cycles of 95°C for 30secs, 60°C for 30s, 72°C for 90secs, and a final extension of 72°C for 10 mins. For arthropods, we amplified a 157bp region of the COI gene using the ZBJ-ArtF1c/R2c primers^5^ (ZBJ-ArtF1c: 5’-AGATATTGGAACWTTATATTTTATTTTTGG-3’; ZBJ-ArtR2c: 5’- WACTAATCAATTWCCAAATCCTCC-3’), and the following thermocycling conditions: 95°C for 15min, followed by 35 cycles of 94°C for 40secs, 40°C for 1min, 72°C for 30secs, and a final extension of 72°C for 10 mins. Finally, for plants and fungi we amplified a 580bp region of the ITS2 region with the S2F/ITS4 primers^6^ (S2F: 5’-ATGCGATACTTGGTGTGAAT-3’; ITS4: 5’-TCCTCCGCTTATTGATATGC-3’). The thermocycling conditions were: 95°C for 10min, followed by 40 cycles of 94°C for 40secs, 49°C for 40secs, 72°C for 40secs, and a final extension of 72°C for 10 mins. All the primers were modified with the CS1/CS2 adaptors. We included six PCR negative controls and the PCR1 reactions were performed in triplicates, except for 12S birds (birT-F/R) which was run only on a subset of samples from the following sites: Coedgwilym Cemetery, Belfast Center, Port Talbot Margam, Scunthorpe Low Santon, Chadwell St Mary, London Marylebone Road, Sheffield Devonshire Green and Auchencorth Moss. We also included a positive control for each region: brown anole (*Anolis sagrei*) for 16S vertebrates and 12S mammals, black guillemot (*Cepphus grylle*) for 12S birds, *Daphnia pulex* for COI and banana (*Musa*) for ITS2. The technical replicates were pooled and transported on ice to the Canadian Centre for DNA Barcoding (CCDB, Guelph, Ontario) for PCR2 (indexing including the M13 tails), cleaning, quantification, normalization, pooling and sequencing on Illumina NextSeq2000 sequencer (2x300 PE cycle run, P1 reagent kit).

Note that we selected the MiMammal-U-F/R (mammals), mam1/2 (vertebrates), ZBJ-ArtF1c/R2c (arthropods), and S2F/ITS4 (plants) primers to capture a broad taxonomic diversity across trophic levels. These primers have previously demonstrated success in amplifying airborne eDNA in previous studies, including air quality monitoring networks^12^. To our knowledge, the birT-F/R (birds) primers had not been previously tested on airborne eDNA samples. However, they were specifically designed for avian environmental DNA metabarcoding, making them a suitable candidate for evaluation in this study.

*Bioinformatic steps and parameters*

Raw reads were first processed by the CCDB. R1 and R2 reads were assembled using the PEAR v0.9.11 program^7^ with the following parameters: statistical method = OES, p-value = 0.01, quality score threshold trimming = 0, maximum assembly length = 999999, minimum assembly length = 50, minimum read size after trimming = 1, maximal ratio of uncalled bases = 1, minimum overlap = 10 bp, and scoring method = scaled score. Cutadapt v3.7^8^ was used to demultiplex assembled reads by sample, remove M13 adapters (-e 0.2 -m 100 -u -17/18), and filter reads (-m 100 -e 0.125 --action=none -g sequence --discard-untrimmed). The full index combinations were searched during the demultiplexing step even if they were not used in our project (“ghost plates”) to estimate assignment errors.

Using cutadapt 4.5, the reads were split by primer pair and the adapters CS1/2 were trimmed using the linked adapters option (-g "CGGTTGGGGTGACCTCGGA;e=0.2...AGTTACCCTAGGGATAACAGC;e=0.2" -m 10). Then, the reads were processed using the DADA2 pipeline v1.26.0^9^ in R 4.4.0^10^ and RStudio^11^. Using the filterAndTrim function, reads were truncated after 157 bp (COI ZBJ-ArtF1c/R2c), 171 bp (12S MiMammal-U-F/R), 90 bp (16S mam1/2), 265 bp (12S birT-F/R) or 230 bp (ITS2 S2F/ITS4), and reads matching against the phiX genome (rm.phix = T), with ambiguous bases (maxN = 0), or with more than 2 expected errors (maxEE = 2) were discarded. All identical sequencing reads were then combined into unique sequences (dereplication) using derepFastq with default parameters. The error rate (learnErrors function) was used to make ASVs with the dada() function. Finally, chimeras were removed using the removeBimeraDenovo function (consensus method).

Taxonomic assignment was kept at the species level only with query cover > 60% and percentage of identity ≥ 97%, and at the order level with percentage of identity < 97% (Appendix S1). In cases where multiple species matched with the same percentage of identity, the closest common ancestor (e.g., genus) was retained. To avoid overinflation of taxonomic assignments at the species level for ITS2 and COI, genus-level identifications were retained for these two regions^12^. For the 16S and two 12S markers we used the online BLAST tool (NCBI full nucleotide collection) to manually assign taxonomy to the ASVs. For the primers with large numbers of ASV (4,389 and 7,550 ASVs for ITS2 and COI, respectively) we combined automated (non-Bayesian taxonomy classifier, sintax command of the usearch v11.0.667 pipeline^13^ and manual (online BLAST tool, NCBI full nucleotide collection) approaches. For COI, we used syntax against the BOLD database. ASVs with a sintax probability at the genus level ≥ 0.60 were checked manually in GenBank (May 2024). For those with a probability at the genus level < 0.60 and family level ≥ 0.60, the ASV was assigned at the family level. If the probability was < 0.60 at the family level, we checked the order level: if the probability was ≥ 0.60, then the ASV was assigned at the order level. This was repeated up to the phylum level: if the phylum level probability was < 0.60, then the ASV was identified as "no match". The same backward strategy was applied to ITS2, with two databases PLANiTS_29_03_2023^14^ and Unite all eukaryotes v10.0^15^.

*Sequencing results*

A total of 118,240,346 raw reads were generated, 117,051,477 were assembled and 97,963,931 were trimmed, filtered and demultiplexed by sample. Following demultiplexing by primer pair in cutadapt and filtering, denoising and chimera removal in DADA2, we obtained 14,381,711 reads and 2,422 ASVs for 12S MiMammal-U-F/R, 24,074,614 reads and 1,496 ASVs for 16S mam1/2, 7,576,673 reads and 1,180 ASVs for 12S birT-F/R, 16,222,647 reads and 7,550 ASVs for ZBJ-ArtF1c/R2c, and 11,813,754 reads and 4,389 ASVs for ITS2 S2F/ITS4 (Table S1). The final dataset, following removal of no match and suspected contaminants, filtering with control “ghost” plates and negative controls, removal of positive control samples and merging of ASVs with identical identifications, had a total of 18,373,005 reads and 1,556 ASVs (details in Table S1) with an average of 1,224,867 ± 486,829 reads and 334 ± 93 ASVs per sampling site (min: 201 at Heigham Holmes, max: 568 at Low Santon).

*References*

10.	R Core Team. R: A language and environment for statistical computing. R Foundation for Statistical Computing (2024).

11.	Posit team. RStudio: Integrated Development Environment for R. Posit Software (2024).

12.	Littlefair, J. E. *et al.* Air-quality networks collect environmental DNA with the potential to measure biodiversity at continental scales. *Current Biology* **33**, R426–R428 (2023).

13.	Edgar, R. SINTAX: a simple non-Bayesian taxonomy classifier for 16S and ITS sequences. *biorxiv* 074161 (2016).

14.	Banchi, E. *et al.* PLANiTS: a curated sequence reference dataset for plant ITS DNA metabarcoding. *Database* **2020**, baz155 (2020).

15.	Abarenkov, K. *et al.* The UNITE database for molecular identification and taxonomic communication of fungi and other eukaryotes: sequences, taxa and classifications reconsidered. *Nucleic Acids Research* **52**, D791–D797 (2024).
