## Appendix S2 for "First national survey of terrestrial biodiversity using airborne eDNA"

**Appendix S2.** Effect of filter storage time at ambient temperature on ASV recovery.

*Methods*

We assessed the effect of storage time at ambient temperature (measured in months between sample collection and storage at -20°C) on the number of ASV recovered. We used the glmmTMB function of the glmmTMB package^1^ to run a GLMM of negative binomial family (nbinom2) with ambient storage time (months), sequencing depth (transformed read count with the scale function) and their interaction as fixed effects. Site and taxonomic group (vertebrate, arthropod, plant, fungi) variables were included as random effects (intercept). The model results were visualized and reported using the plot_model and tab_model functions of the sjplot package^2^, and the image, contour and persp functions of the rsm package^3^. The analysis was performed on i) all taxa, ii) taxa identified at the species level only, iii) taxa identified at the genus level only, iv) taxa identified at the family level only, and v) taxa identified at the order level only. Note that at the species level, we performed a simpler model (GLM with negative binomial family) because: 1) there was only one taxonomic group (only vertebrates were identified at the species level) so it was not included in the model, and 2) the random effect “site” was not contributing additional explanatory power to the model with null between-group variance of the random effect.

*Results*

We observed a negative effect of ambient storage time, regardless of the taxonomic level, and a positive effect of read count at all levels but family (Appendix 2 Figure 1 and Table below). The negative IRR of the interaction between ambient storage time and read counts at the species level indicates that the longer the ambient storage time, the less “beneficial” the total read count becomes for species richness (Appendix 2 Figure 2).

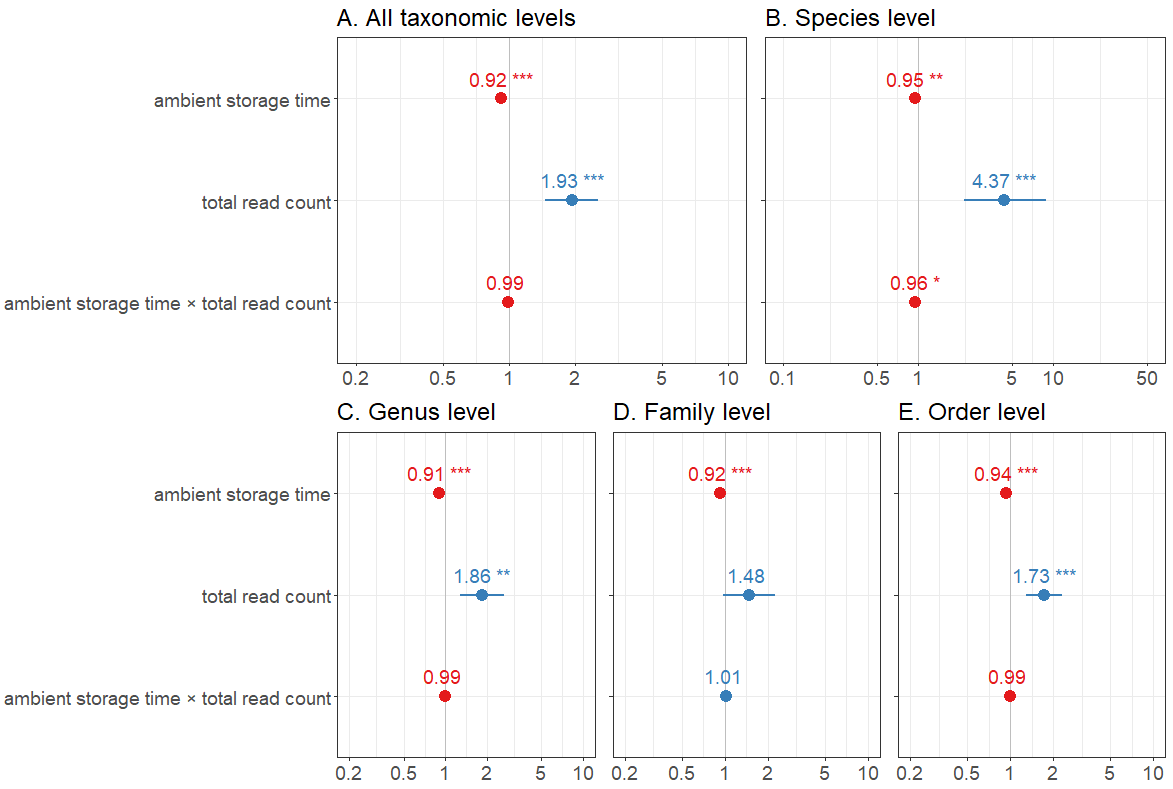

**Appendix 2 Figure 1.** Incidence rate ratios (IRR) and standard errors for each predictor (ambient storage time, total read count and their interaction, for A) ASV identified at any taxonomic level, ii) ASV identified at the species level only, iii) ASV identified at the genus level only, iv) ASV identified at the family level only, and v) ASV identified at the order level only. Predictors with ICC < 1 (left of the grey line; red) have a negative effect, while those >1 (on the right; blue) have a positive effect. The significance (p-value) is indicated by stars.

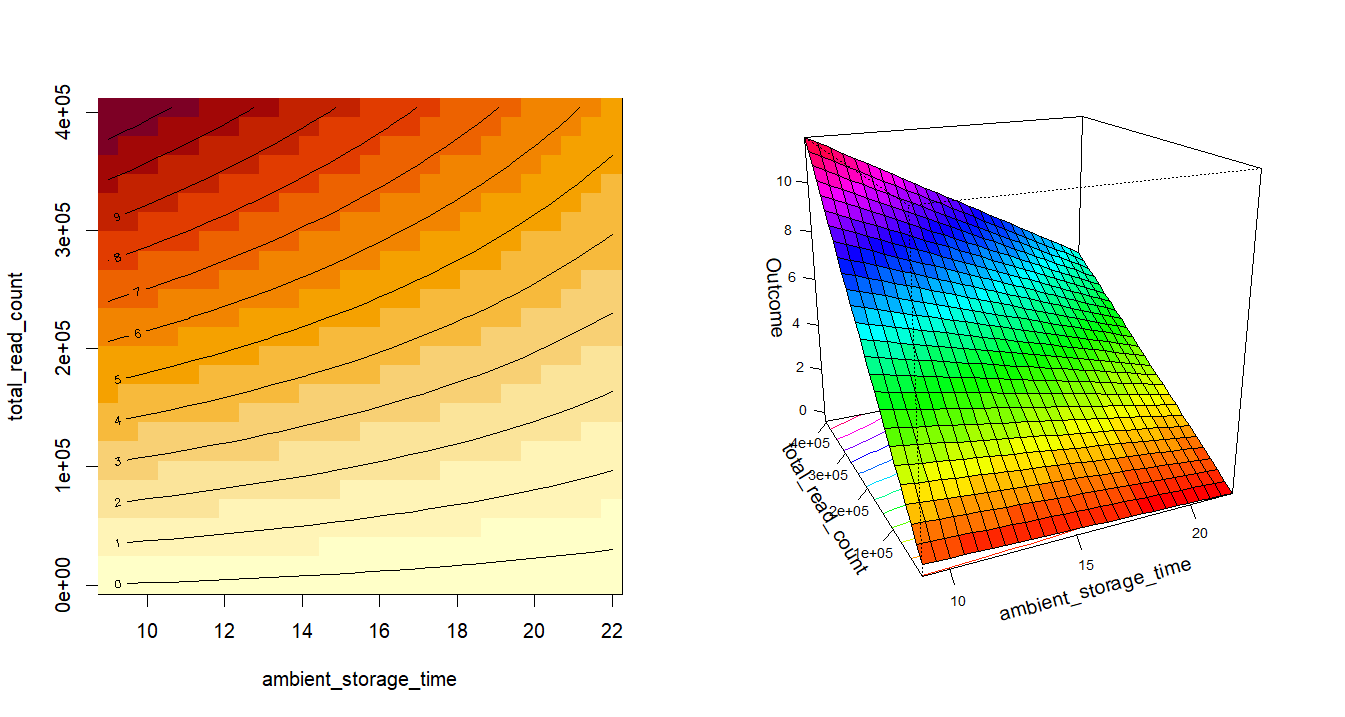

**Appendix 2 Figure 2.** Interaction plots representing the negative interaction between the two predictor variables at the species level: ambient storage time (months) and read counts. Each contour line of the left plot corresponds to a specific level of predicted species richness (e.g. along the line labeled "4" the predicted species richness is 4 for the specific combinations of ambient storage time and total read count that lie along that line), with warmer colors (red) representing higher values of predicted species richness. The right plot is a 3D representation of the outcome (predicted species richness, untransformed estimate) variation across the range of values of the read count and ambient storage time variables. Both plots demonstrate that species richness declines with storage time and reduction in read counts.

**Appendix 2 Table.** Output of the models investigating the effect of ambient storage time of the filters (‘time’), sequencing depth (‘read’) and their interaction on the number of ASV identified at any taxonomic levels, species only, genus only, family only and order only, respectively.

|  | **All taxonomic levels (negative binomial GLMM)** | | | | **Species level (negative binomial GLM)** | | | | **Genus level (negative binomial GLMM)** | | | | **Family level (negative binomial GLMM)** | | | | **Order level (negative binomial GLMM)** | | | |
| --- | --- | --- | --- | --- | --- | --- | --- | --- | --- | --- | --- | --- | --- | --- | --- | --- | --- | --- | --- | --- |
| **Predictors** | **Estimate** | **Std. Error** | **z value** | **p** | **Estimate** | **Std. Error** | **z value** | **p** | **Estimate** | **Std. Error** | **z value** | **p** | **Estimate** | **Std. Error** | **z value** | **p** | **Estimate** | **Std. Error** | **z value** | **p** |
| (Intercept) | 3.057 | 0.374 | 8.178 | **2.88e-16** | 0.908 | 0.260 | 3.494 | **0.0004** | 2.599 | 0.504 | 5.160 | **2.47e-07** | 0.239 | 0.839 | 0.285 | 0.776 | 0.723 | 0.314 | 2.302 | **0.021** |
| time | -0.080 | 0.007 | -11.122 | **< 2e-16** | -0.051 | 0.017 | -3.000 | **0.003** | -0.097 | 0.010 | -9.864 | **< 2e-16** | -0.084 | 0.013 | -6.331 | **2.43e-10** | -0.062 | 0.011 | -5.887 | **3.94e-09** |
| read | 0.660 | 0.143 | 4.621 | **3.82e-06** | 1.475 | 0.236 | 6.247 | **4.18e-10** | 0.619 | 0.189 | 3.270 | **0.001** | 0.395 | 0.212 | 1.859 | 0.063 | 0.548 | 0.148 | 3.696 | **0.0002** |
| time × read | -0.013 | 0.010 | -1.326 | 0.185 | -0.041 | 0.016 | -2.517 | **0.012** | -0.006 | 0.013 | -0.492 | 0.623 | 0.011 | 0.014 | 0.800 | 0.424 | -0.009 | 0.010 | -0.863 | 0.388 |
| **Random Effects** | **Type** | **Variance** | **Std. Dev** |  | **Type** | **Variance** | **Std. Dev** |  | **Type** | **Variance** | **Std. Dev** | | **Type** | **Variance** | **Std. Dev** |  | **Type** | **Variance** | **Std. Dev** |  |
| Supergroup | Intercept | 0.618 | 0.786 |  | N/A | | | | Intercept | 0.073 | 0.269 |  | Intercept | 2.926 | 1.711 |  | Intercept | 0.341 | 0.584 |  |
| site | Intercept | 0.031 | 0.178 |  |  |  |  |  | Intercept | 1.117 | 1.057 |  | Intercept | 0.075 | 0.274 |  | Intercept | 0.022 | 0.150 |  |
| Marginal / Conditional R^2^ | 0.223 / 0.641 | | | | 0.309 (R² Nagelkerke) | | | | 0.186 / 0.680 | | | | 0.100 / 0.786 | | | | 0.165 / 0.400 | | | |

*References*

1.	Brooks, M. E. *et al.* glmmTMB Balances Speed and Flexibility Among Packages for Zero-inflated Generalized Linear Mixed Modeling. *The R Journal* 9, 378–400 (2017).

2.	Lüdecke, D. sjPlot: Data Visualization for Statistics in Social Science. R package version 2.8.16. (2024)

3.	Russell V. Lenth. Response-Surface Methods in R, Using rsm. *Journal of Statistical Software*, 32(7), 1-17. (2009)
