## Appendix S3 for "First national survey of terrestrial biodiversity using airborne eDNA"

**Appendix S3. Comparison of iNaturalist and airborne eDNA datasets within an 18.6 km radius**

Similarly to eBird, the iNaturalist dataset comprised more taxa than airborne eDNA, with 2,942 taxa from 17,078 observations compared to 1,227 taxa from 185 samples, respectively. However, the overlap in shared species-site detections between the two datasets was lower (7.4%) than when comparing eBird and eDNA with a higher overall detection rate by iNaturalist than airborne eDNA (McNear’s Chi-squared 𝜒 = 6005.8, p < 0.001), with 43.0% found only in the iNaturalist dataset, and 49.9% found only in the eDNA dataset (**Appendix 3 Figure 1**). Airborne eDNA detected some taxa not recovered with iNaturalist (Branchiopoda, Copepoda, Nematoda, Rotifera, Tardigrada, Chytriomycota, Chlorophyta, Charophyta and Cryptista) (**Appendix 3 Figure 1**, Table S4). Similarly, iNaturalist detected some taxa not recovered with eDNA (Reptilia, Mollusca, Pycnogonida, Cnidaria, Porifera, Echinodermata, platyhelminthes, tunicates, Chromista, Protozoa, Rodophyta, Mucoromycota, Polypodiopsida, Ceratophyllales, Eumagnoliids, Nymphaeales and Lycophytes) (**Appendix 3 Figure 1**, Table S4).

Airborne eDNA recovered more detections of Ascomycota (𝜒 = 26, p < 0.001), Chlorophyta (𝜒 = 101.4, p < 0.001) and mammals (𝜒= 5.7, p = 0.016). iNaturalist recovered more detections of Basidiomycota (𝜒 = 630.9, p < 0.001), Bryophyta (𝜒 = 164.5, p < 0.001), Monocots (𝜒 = 299.6, p < 0.001), Eudicots (𝜒 = 2231.5, p = 0), Marchantiophyta (𝜒 = 47.2, p < 0.001), Insecta (𝜒 = 2424.8, p = 0), Arachnida (𝜒 = 141.5, p < 0.001), Malacostraca (𝜒 = 49.9, p < 0.001), Diplopoda (𝜒 = 18.2, p < 0.001), Fish (𝜒 = 54.8, p < 0.001), Bird (𝜒 = 615.8, p < 0.001) and Amphibian (𝜒 = 45.2, p < 0.001). No significant difference was observed for Bryozoa (p = 1.000), Ostracoda (p = 1.000), Gymnosperm (p = 0.371), Thecostraca (p = 0.617), Entognatha (p = 0.065) and Annelida (p = 0.742).

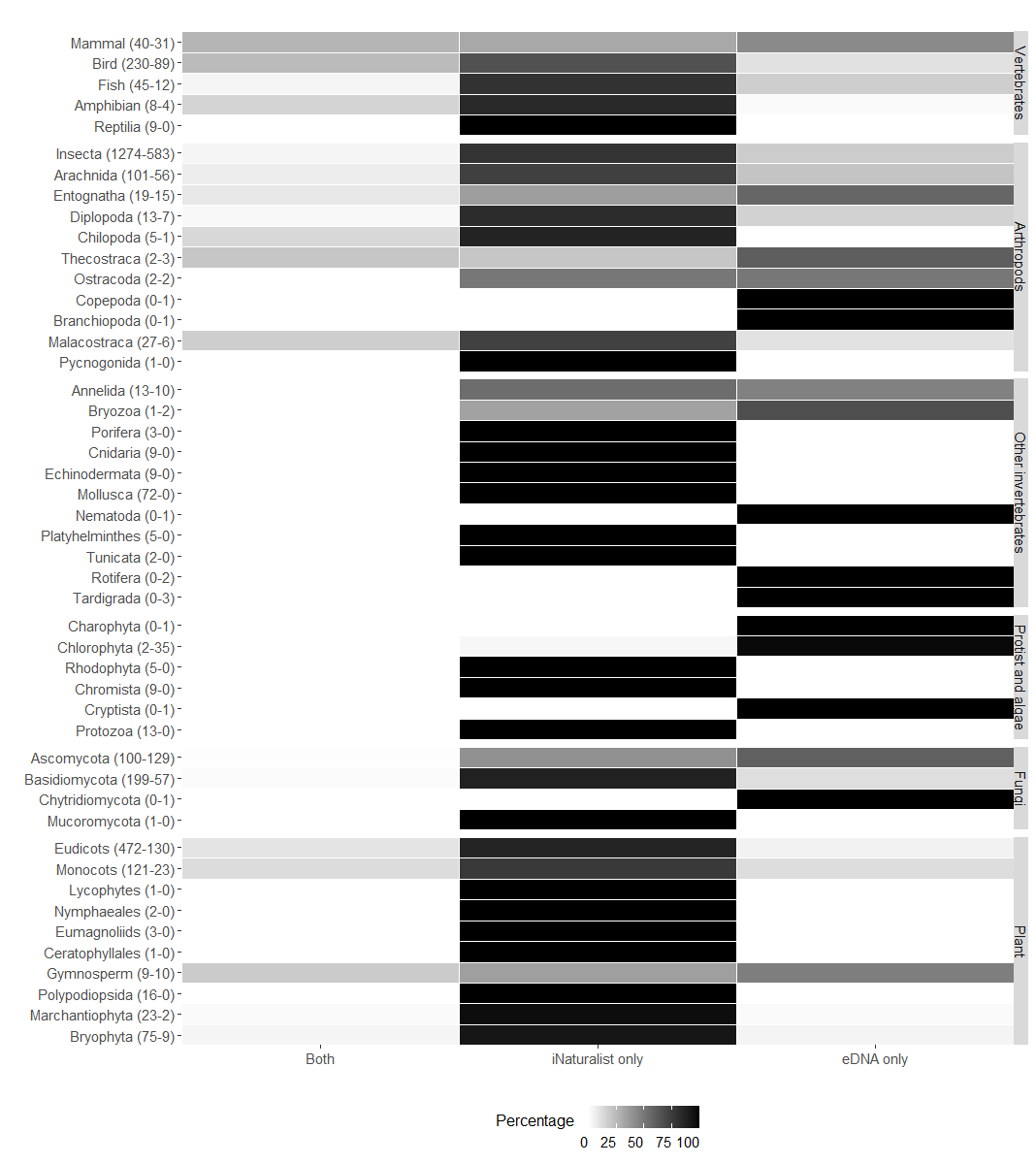

**Appendix 3 Figure 1.** Detection source percentage per taxonomic group (vertebrates, arthropods, other invertebrates, fungi, plants, protists and algae) in a 18.6km radius around the sampling sites. “Both” = percentage of detections in both the iNaturalist and eDNA datasets, “iNaturalist only” = percentage of detections in the iNaturalist dataset only, and “eDNA only” = percentage of detections in the eDNA dataset only. The numbers of taxa in the iNaturalist checklists (total of 2,942 taxa from 17,078 observations) and the eDNA dataset (1,227 taxa from 185 samples) are indicated between brackets for each taxonomic group.
