## Supplementary material for "First national survey of terrestrial biodiversity using airborne eDNA": Figure S1

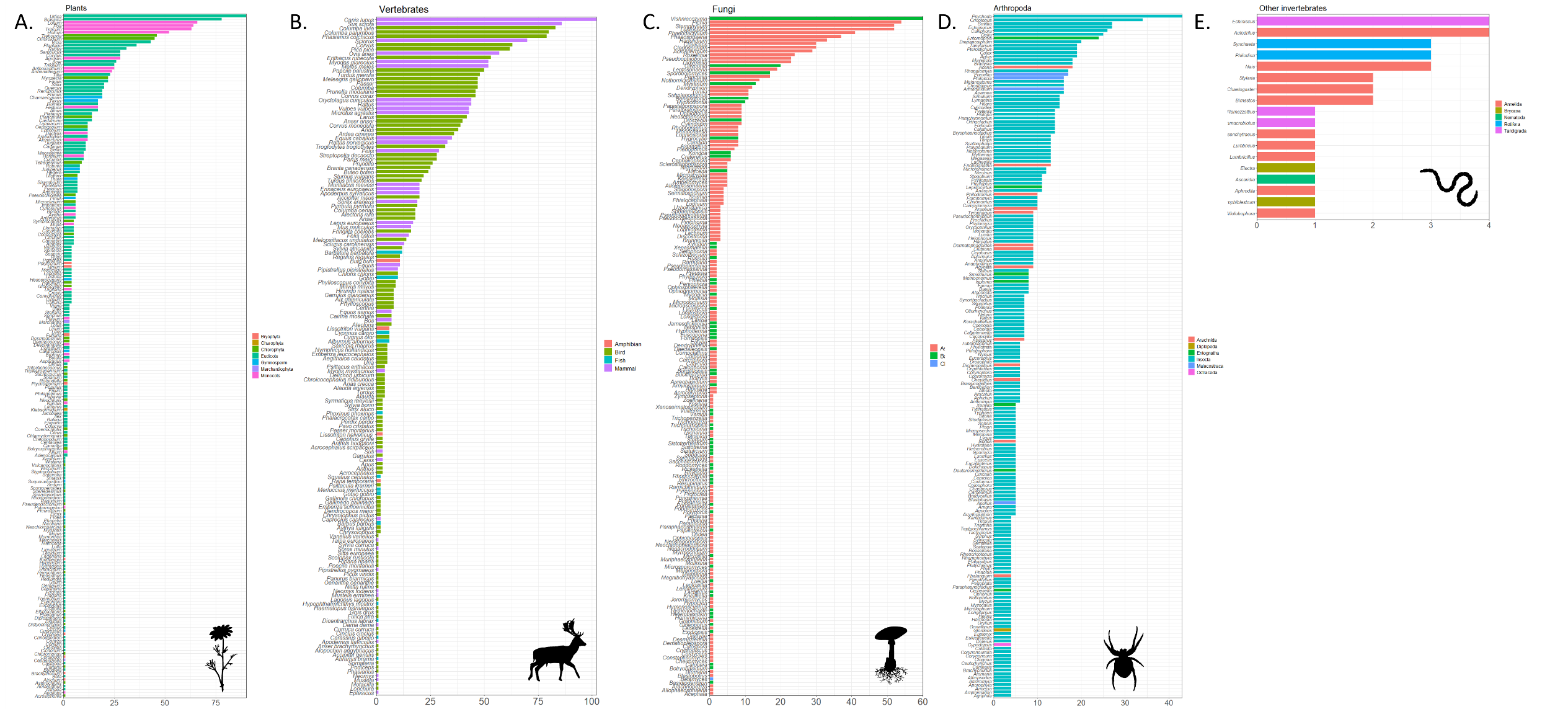

**Figure S1**. Detection counts (i.e. positive samples) of each A) plant (genus), B) vertebrate (genus and species), C) fungi (genus), D) arthropod (genus) and E) non-arthropod invertebrate (genus) taxa in the airborne eDNA dataset, sorted from the most to the least detected in our dataset. For more clarity, arthropod taxa with less than three detections are not represented here.
