## Supplementary material for "First national survey of terrestrial biodiversity using airborne eDNA": Figure S2

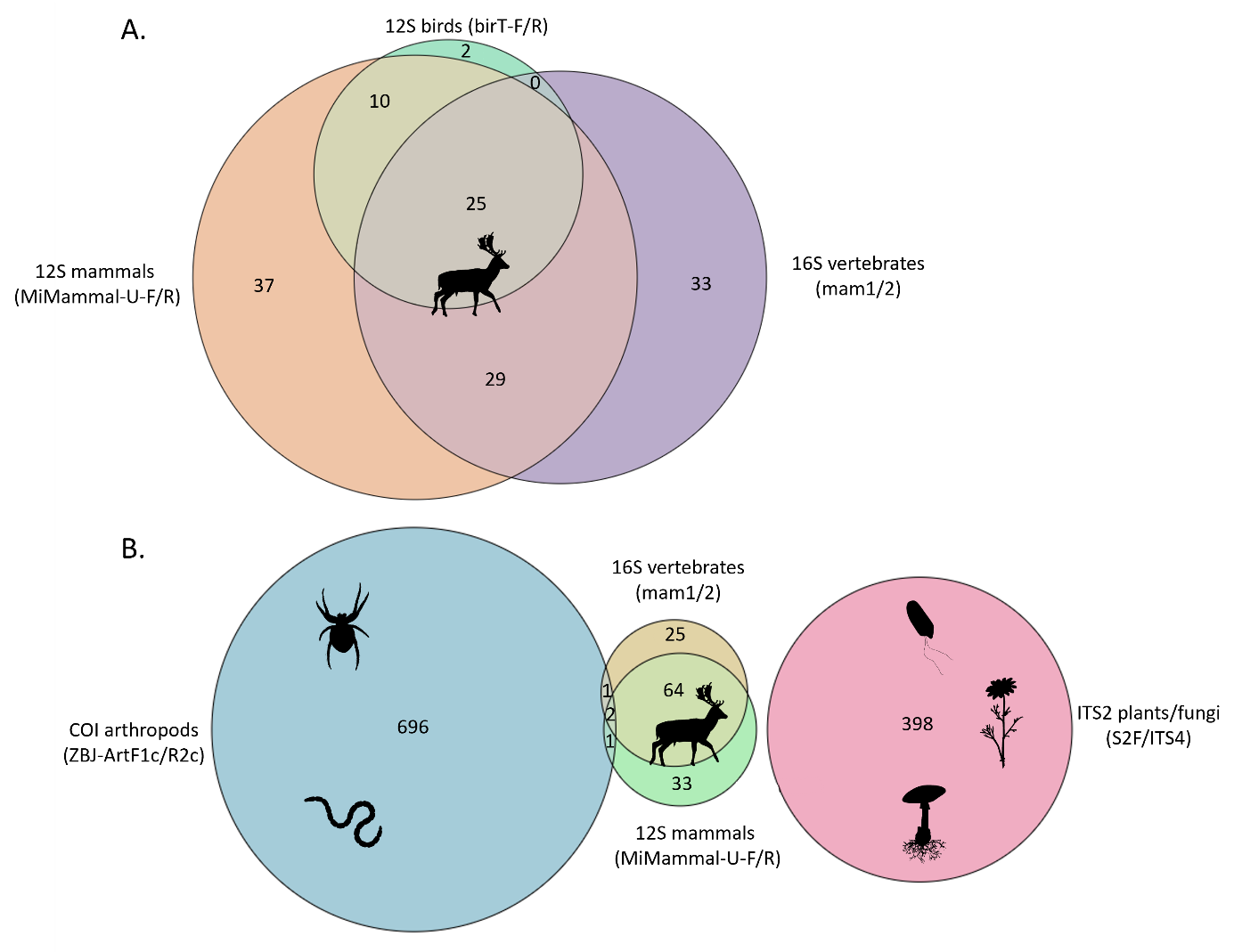

**Figure S2.** Overlap of A) vertebrate taxa at the species level (birds, mammals, fish, amphibians) based on two markers in three primer pairs: 12S birds (birT-F/R), 12S mammals (MiMammal-U-F/R), 16S vertebrates (mam1/2), and B) all taxa at the genus level (arthropods, other invertebrates, vertebrates, fungi, plants and protists) based on four markers in five primer pairs: 12S birds (birT-F/R), 12S mammals (MiMammal-U-F/R), 16S vertebrates (mam1/2), COI arthropods (ZBJ-ArtF1c/R2c), ITS2 plants/fungi (S2F/ITS4). Circle size is proportional to the number of taxa. Note that 12S birds (birT-F/R) completely overlaps with the other primer pair at the genus level.
